# Secondary Metabolites from *Munnozia hastifolia* (Asteraceae)

**DOI:** 10.64898/2026.10.16.758089

**Authors:** Rosana Alarcón, Alejandro Alarcón, Abel Flores, Diego Germán Gutiérrez, Nélida Bayón

**Author notes:** Contactos* | *Contacts:* Rosana Alarcón. *E-mail address:.

## Abstract

A phytochemical investigation of the flowers of *Munnozia hastifolia* led to the isolation of four known compounds tulirinol **1**, dihydrochrysanolide **2**, quercetin **3** and luteolin **4**, while the leaves afforded lupeol **5**, α-amyrin **6** and stigmasterol **7**. The structures were determinated using several spectroscopic data. This is the first report about chemical composition of *M. hastifolia*.

## INTRODUCTION

The genus *Munnozia* (Asteraceae, tribe Liabeae) contains about 40 species, occurring from Colombia to northwestern Argentina (Dillon et al., 2009; Funk & Robinson 2001; Funk et al., 2012; Robinson 1983) of which only *Munnozia hastifolia* (Poepp.) H. Rob. and Brettell grows in Argentina (Gutiérrez and Luna, 2013). In Perú, it is traditionally known as “alas de murciélago” and several medicinal properties are attributed to this plant (Luziatelli et al., 2010).

Previous studies have indicated that plants of the genus *Munnozia* contain sesquiterpene lactones, triterpenes, polyacetylenes and thiophene derivatives (Bohlmann et al., 1977; Bohlmann & Grenz, 1979; Fournet et al., 1993). Leaf extracts of *Munnozia fournetii* and *Munnozia maronii* have shown antileishmanial activity (Fournet et al., 1993; Rocha et al., 2005), but bark extracts of *M. hastifolia* are not active (Valadeau et al., 2009). Recently, studies have been conducted to assess the topical efficacy of *M. hastifolia* ointments with promising results (Pongo Alcalá & Herrera Flores, 2015).

At the best of our knowledge, there are not previous reports about *M. hastifolia* chemical composition, so this is the first contribution to its chemical analysis. The main goal of this research was to study the compounds of *M. hastifolia* grown on northwestern Argentina.

## MATERIAL AND METHODS

### General

NMR spectra were recorded on a Bruker Avance 400 (^1^H at 400 MHz and ^13^C at 100 MHz) spectrometer with TMS as internal reference. HRMS were performed on a Bruker micrOTOF-QII spectrometer. IR spectra were taken on an IR-FT Bruker model IFS-88 spectrometer. CC were performed on silica-gel 230-400 mesh (Fluka), RPCC on C-18 silica gel (Merck), TLC was carried out on precoated Silica gel 60 F_254_ plates (Fluka). Detection was achieved by UV light and spraying with 10 % vanillin in EtOH followed by heating.

#### Plant material and extraction

Aerial parts (leaves, flowers) of *M. hastifolia* were collected in August 2010, in the province of Jujuy, Argentina. A voucher specimen (MCNS 10379) is on deposit at the Museo de la Facultad de Ciencias Naturales, Universidad Nacional de Salta, Salta, Argentina.

#### Isolation and identification

The flowers (150 g) of *M. hastifolia* were macerated in EtOH for 7 days at room temperature. The organic solution was distilled under reduced pressure at 40°C to obtain 32.0 g of crude extract. This extract was defatted by dissolving in MeOH and precipitation at −10 °C. After removal by filtration of the abundant waxy precipitate, the filtrate was evaporated to afford 12.3 g of a gummy residue, which was divided into 2 fractions by flash column chromatography on silica gel C-18 (70 g) (7 × 15 cm), eluting with MeOH:H_2_O (7:3, 500 mL) and MeOH (500 mL).

The first fraction (5.5 g) was subjected to flash column chromatography on silica gel using hexane-EtOAc mixtures of increasing polarity to yield 6 fractions. *Fraction 3* (25 mg) eluted with hexane-EtOAc (6:4) was purified by preparative TLC (hexane-EtOAc 5:1, runs x 2) to afford 1.5 mg of **1** (Rf= 0.3). *Fraction 4* (30 mg) eluted with hexane-EtOAc (4:6) gave 2.2 mg of **2**, after purification by preparative TLC (CH_2_Cl_2_-Me_2_CO 2.9:0.1, runs x 2, Rf= 0.7). *Fraction 5* (50 mg) eluted with hexane-EtOAc (2:8) was chromatographed by flash CC on silica gel C-18 using MeOH-H_2_O (8:2) to yield **3** (3.5 mg), and a mixture that was submitted to preparative TLC with hexane-Et_2_O-EtOAc (4:1:3) to afford **4** (2.0 mg). Air-dried leaves (80 g) were macerated with EtOH for 1 week at room temperature. After filtration, the solvent was removed *in vacuo* to yield a residue (3.5 g), which was submitted to flash column chromatography on silica gel C-18, using MeOH-H_2_O 9:1 (*Fraction 1*) and MeOH (*Fraction 2). Fraction 1* (1.7 g) was first purified by column chromatography on silica gel using mixtures (100 mL each) of n-hexane-Et_2_O of increasing polarity (10 %); fractions of 10 mL were collected. Fractions eluted with n-hexane-Et_2_O (8:2) contained a mixture of **5** and **6** (10 mg). Fractions eluted with n-hexane-Et_2_O (7:3) gave stigmasterol **7** (20 mg).

### Characterization of the compounds

#### Tulirinol (1)

Amorphous solid; IR ν (KBr) ν_max_ 3432, 2929, 2852, 1767, 1732, 1447, 1245, 1032; For ^1^H NMR and ^13^C NMR data, see Table 1; HRESIMS: 329.13811 [M+Na]^+^ (calcd for C_17_H_22_ O_5_Na, 329.13701); EI MS (70 eV) m/z: [M+Na] ^+^ 329 (100), 294 (12), 229 (12), 101 (67).

**Table 1.** ^1^H (400 MHz), ^13^C (100 MHz) NMR data (δ in ppm), HMBC and NOESY correlations of compound **1** (CDCl_3_).

| Position | $\delta$ $^{13}\text{C}$ | $\delta$ $^1\text{H}$ | HMBC<br>($^1\text{H} \rightarrow ^{13}\text{C}$) | NOESY |
| --- | --- | --- | --- | --- |
| **1** | 66.8 | 4.40 dd<br>(11.1; 5.3) | C-9, C-14 | H-8 |
| **4** | 27.3 | 2.00<br>1.73 m | C-1, C-10 |  |
| **3** | 35.3 | 1.89 - 2.30 m | C-2, C-4, C-5, C-15 |  |
| **4** | 137.9 | - | - |  |
| **5** | 126.0 | 4.86 br d<br>(10.7) | C-3, C-15 |  |
| **6** | 73.1 | 5.39 t<br>(10.2) | C-4, C-7, C-1' | H-15, H-8, H-13 |
| **7** | 49.2 | 3.03 m | C-11 |  |
| **8** | 74.1 | 4.62 t<br>(9.5) | C-6, C-7, C-9 | H-1, H-6 |
| **9** | 126.4 | 5.33 br d<br>(10.0) | C-1, C-14 | H-14 |
| **10** | 142.8 | - | - |  |
| **11** | 136.9 | - | - |  |
| **12** | 169.3 | - | - |  |
| **13** | 123.0 | 5.75 d<br>(3.0)<br>6.28 d<br>(3.4) | C-7, C-11, C-12 | H-6 |
| **14** | 16.8 | 1.83 s | C-1, C-9, C-10 | H-9 |
| **15** | 15.6 | 1.90 s | C-3, C-4, C-5 | H-6 |
| **OAc** | 169.9 | - | - |  |
|  | 21.1 | 2.10 s<br>(6.5) | C-1' |  |

Coupling constant values (in parentheses) are in Hz.

#### Dihydrochrysanolide (2)

Amorphous solid; IR ν (KBr) ν_max_ 3426, 2922, 2850, 1735, 1647, 1438, 1239. For ^1^H NMR and ^13^C NMR data, see Table 2; HRESIMS: 329.13783 [M+Na]^+^ (calcd for C_17_H_22_ O_5_Na, 329.13594); EI MS (70 eV) m/z: [M+Na] ^+^ 329 (100), 291 (1), 219 (11), 101 (9).

**Table 2.** ^1^H (400 MHz), ^13^C (100 MHz) NMR data (δ in ppm) and HMBC correlations of compound **2**.

| Position | δ <sup>13</sup>C<sup>a</sup> | δ <sup>13</sup>C<sup>b</sup> | δ <sup>1</sup>H<sup>a</sup> | δ <sup>1</sup>H<sup>b</sup> | HMBC<sup>a</sup><br>(<sup>1</sup>H → <sup>13</sup>C) |
| --- | --- | --- | --- | --- | --- |
| **1** | 69.0 | 70.2 | 3.90 dd<br>(6.7; 14.0) | 3.85 m | C-10, C-14 |
| **4** | 28.7 | 30.8 | 1.99- 1.96 m | 2.02-2.10 m | C-3 |
| **3** | 34.0 | 34.1 | 2.00- 2.20 m | 2.25-2.10 m | - |
| **4** | 137.8 | 138.2 | - | - | - |
| **5** | 127.1 | 127.0 | 4.96 m | 4.94 m | - |
| **6** | 72.3 | 72.5 | 5.36 m | 5.20 m | C-1' |
| **7** | 48.8 | 48.8 | 3.15 m | 3.01 m | - |
| **8** | 78.3 | 78.7 | 4.16 m | 4.02 m | - |
| **9** | 41.7 | 41.9 | 2.80-2.43 m | 2.95-2.40 m | C-10 |
| **10** | 147.1 | 146.5 | - | - | - |
| **11** | 137.3 | 136.0 | - | - | - |
| **12** | 168.8 | 169.5 | - | - | - |
| **13** | 122.5 | 124.7 | 5.86 dd<br>(0.9; 2.7)<br>6.13 dd<br>(0.9; 3.1) | 5.84 d<br>(2.1)<br>6.35 d<br>(2.9) | C-7, C-11, C-12 |
| **14** | 113.1 | 114.6 | 5.12 s, br<br>5.14 s, br | 5.11 s, br<br>5.16 s, br | C-1, C-9, C-10 |
| **15** | 16.5 | 17.2 | 1.83 s | 1.80 s | C-3, C-4, C-5 |
| **OAc** | 169.2<br>20.1 | 169.7<br>20.9 | -<br>2.06 s | -<br>2.08 s | -<br>C-1' |

Coupling constant values (in parentheses) are in Hz.
<sup>a</sup>In Me<sub>2</sub>CO-d<sub>6</sub><sup>b</sup>In CDCl<sub>3</sub>

#### Quercetin (3)

Data unreported ^13^C NMR (100 MHz, Me_2_CO-d_6_): 146.5 (C-2), 137.2 (C-3), 175.8 (C-4), 161.6 (C-5), 98.1 (C-6), 164.1 (C-7), 93.8 (C-8), 157.1 (C-9), 103.3 (C-10), 122.8 (C-1`), 114.8 (C-2`), 145.9 (C-3`), 147.3 (C-4`), 115.6 (C-5`), 120.7 (C-6`).

## RESULTS AND DISCUSSION

The EtOH extract of the flower heads of *M. hastifolia* afforded four known compounds tulirinol **1** (Doskotch et al., 1980), dihydrochrysanolide **2** (Doskotch et al., 1983; El Feraly et al., 1983; Lee et al., 2002), quercetin **3** (Alarcón et al., 2007) and luteolin **4** (Özgen et al., 2011). The structures of these compounds were determined mainly on the basis of NMR experiments and by comparing their spectral properties with those reported in literature.

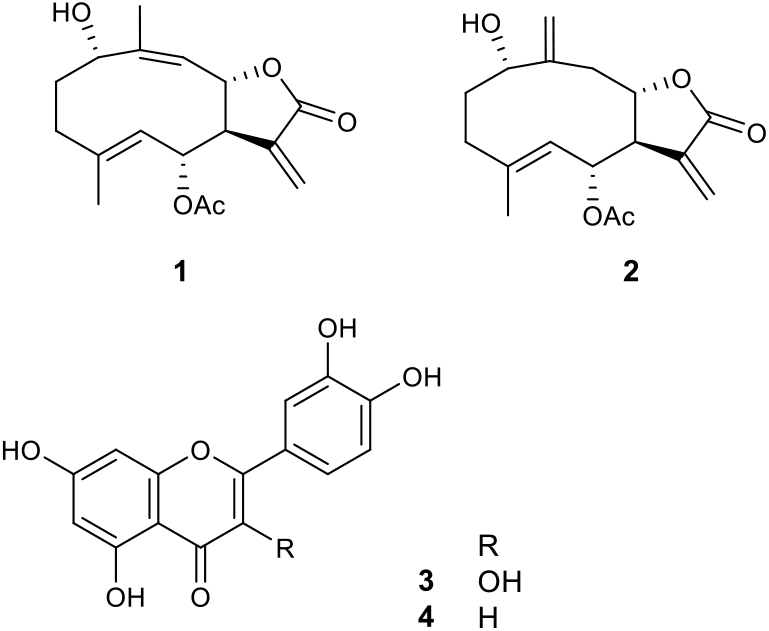

The combined inspection of 2D NMR spectra (COSY, HSQC, HMBC) led to the unambiguous of all ^1^H and ^13^ C NMR resonances for compounds **1** (Table 1) and **2** (Table 2).

The phytochemical analysis of the ethanolic extract obtained from the leaves of *M. hastifolia* revealed the presence of lupeol **5**, α-amyrin **6** y stigmasterol **7**.

## CONCLUSIONS

Compounds **1**-**7** are now reported for the first time in the genus *Munnozia*. Also, in this investigation, it is reported the second *Munnozia* species that synthetizes germacranolides with the lactone between C-7 and C-8 (Bohlmann & Grenz, 1979).

Our results shown that *M. hastifolia* is a source of sesquiterpene lactones with remarkable biological activities (Dosktoch et al., 1980; Lee et al., 2002; Lee et al., 2003a; Lee et al., 2003b).

This is the first report of the presence of tulirinol (**1**) and dihydrochrysanolide (**2**) in a species of *Munnozia* and provides a basis for future studies on bioprospecting *M. hastifolia*.

## ACKNOWLEDGEMENTS

Financial support for this work by the Consejo de Investigación de la Universidad Nacional de Salta is gratefully acknowledged.

